# Differences in neuroinflammation in the olfactory bulb between D614G, Delta and Omicron BA.1 SARS-CoV-2 variants in the hamster model

**DOI:** 10.1101/2022.03.24.485596

**Authors:** Lisa Bauer, Melanie Rissmann, Feline F. W. Benavides, Lonneke Leijten, Lineke Begeman, Edwin Veldhuis Kroeze, Peter van Run, Marion P. G. Koopmans, Barry Rockx, Debby van Riel

## Abstract

Severe acute respiratory syndrome coronavirus 2 (SARS-CoV-2) infection is associated with various neurological complications. SARS-CoV-2 infection induces neuroinflammation in the central nervous system (CNS), whereat the olfactory bulb seems to be involved most frequently. Here we show differences in the neuroinvasiveness and neurovirulence among SARS-CoV-2 variants in the hamster model five days post inoculation. Replication in the olfactory mucosa was observed in all hamsters, but most prominent in D614 inoculated hamsters. We observed neuroinvasion into the CNS via the olfactory nerve in D614G-, but not Delta (B.1.617.2)- or Omicron BA.1 (B.1.1.529) inoculated hamsters. Neuroinvasion was associated with neuroinflammation in the olfactory bulb of hamsters inoculated with D614G but hardly in Delta or Omicron BA.1. Altogether, this indicates that there are differences in the neuroinvasive and neurovirulent potential among SARS-CoV-2 variants in the acute phase of the infection in the hamster model.

## Introduction

The severe acute respiratory syndrome coronavirus 2 (SARS-CoV-2) causing Coronavirus disease 2019 (COVID-19) is associated with wide range of neurological complications in the acute phase and post-acute stage. In the acute stage these include loss of smell (anosmia), headache, fatigue, seizures, confusion and cerebrovascular injuries (Misra et al., 2021; Nasserie et al., 2021). Although the frequency varies among studies, a substantial amount of patients suffer from neurological manifestations in the acute phase of disease (Chou et al., 2021; Spudich and Nath, 2022). Neuropsychiatric maladies such as depression, anxiety and cognitive problems may also persist in the post-acute stage, a clinical picture often referred to as Long Covid (Blomberg et al., 2021; Nolen et al., 2022; Tang et al., 2022; Varatharaj et al., 2020). The underlying mechanisms of the neurological symptoms are barely understood, but given the numerous complications, it is plausible that multiple mechanisms contribute to the neurological pathophysiology.

Currently it is unknown if (emerging) SARS-CoV-2 variants differ in their neuroinvasiveness, neurotropism and neurovirulence. Most studies modelling neurological complications focused on the D614G strain *in vivo* (Frere et al., 2022; de Melo et al., 2021; Wenzel et al., 2021; Zazhytska et al., 2022). Evidence suggests that D614G is neuroinvasive, in at least a subset of patients and experimentally inoculated animals (Meinhardt et al., 2021; de Melo et al., 2021). Although SARS-CoV-2 might enter the central nervous system (CNS) via different routes, evidence suggests that the olfactory nerve is an important route of entry into the CNS (Bauer et al., 2022). The olfactory nerve connects the olfactory mucosa directly with the olfactory bulb in the brain and thus represents a shortcut between the nasal cavity and the brain (Riel et al., 2015). In the CNS, the neurotropism of SARS-CoV-2 appears to be restricted to a subset of permissive CNS cells based on *in vivo* (Frere et al., 2022; Meinhardt et al., 2021; de Melo et al., 2021; Sia et al., 2020; Zazhytska et al., 2022) and *in vitro* (Bauer et al., 2021; Bullen et al., 2020; Jacob et al., 2020; McMahon et al., 2021; Pellegrini et al., 2020; Ramani et al., 2020; Wang et al., 2021; Zhang et al., 2020) studies. However, SARS-CoV-2 replication in the different cell types of the CNS is often inefficient and abortive (Bauer et al., 2021; Bullen et al., 2020; Frere et al., 2022; McMahon et al., 2021; Ramani et al., 2020; Wang et al., 2021; Zazhytska et al., 2022; Zhang et al., 2020) with the exception of choroid plexus epithelial cells *in vitro* (Jacob et al., 2020; Pellegrini et al., 2020). Despite this inefficient replication, SARS-CoV-2 is neurovirulent and triggers neuroinflammatory responses in different anatomical parts of the brain (Bauer et al., 2021; Frere et al., 2022; Zazhytska et al., 2022).

In order to understand if SARS-CoV-2 variants differ in their neuroinvasiveness, neurotropism and neurovirulence, we studied this in Syrian golden hamsters. We compared hamsters inoculated with D614G, Delta (B.1.617.2) or Omicron BA.1 (B.1.1.529) in their potential for CNS invasion, and the associated antiviral, neuroinflammation and microglial activation in the olfactory bulb, the cerebral cortex and the cerebellum.

## Results

### Neuroinvasiveness of D614G, Delta and Omicron BA.1

Nasal turbinates (containing olfactory mucosa), olfactory bulb, cerebral cortex and cerebellum were collected from hamsters five days post intranasal inoculation with SARS-CoV-2 variant D614G, Delta or Omicron BA.1 five days post inoculation (Figure S1).

No significant difference between the viral titers in the nasal turbinates was observed, although there was a trend of higher titers in hamsters infected with the Delta variant, compared to D614G and Omicron BA.1 (Figure 1A). There was significantly more viral RNA in the nasal turbinates of hamsters inoculated with Delta than in hamsters inoculated with D614G (Fig 1B). In the olfactory epithelium, SARS-CoV-2 antigen was most abundantly found in the D614G hamsters compared to Delta and Omicron BA.1 infected hamsters (Figure 1C and Table S1). In the nasal turbinates, D614G inoculated hamsters showed the most histopathological evidence of inflammation demonstrated by multifocal mild to moderate attenuation of the olfactory epithelium colocalized with positive cytoplasmic immunohistochemical staining for SARS-CoV-2 antigen (Table S1). Within the affected olfactory epithelium and lamina propria predominantly neutrophilic infiltrates were present, whereas the associated turbinates’ lumina contained substantial mucopurulent exudates. As reported by *Rissmann et al.* (Rissmann et al., 2022), the hamsters infected with Delta showed more inflammation compared to Omicron BA.1 infected hamsters, which showed the least inflammatory lesions within the olfactory mucosa.

**Figure 1.**
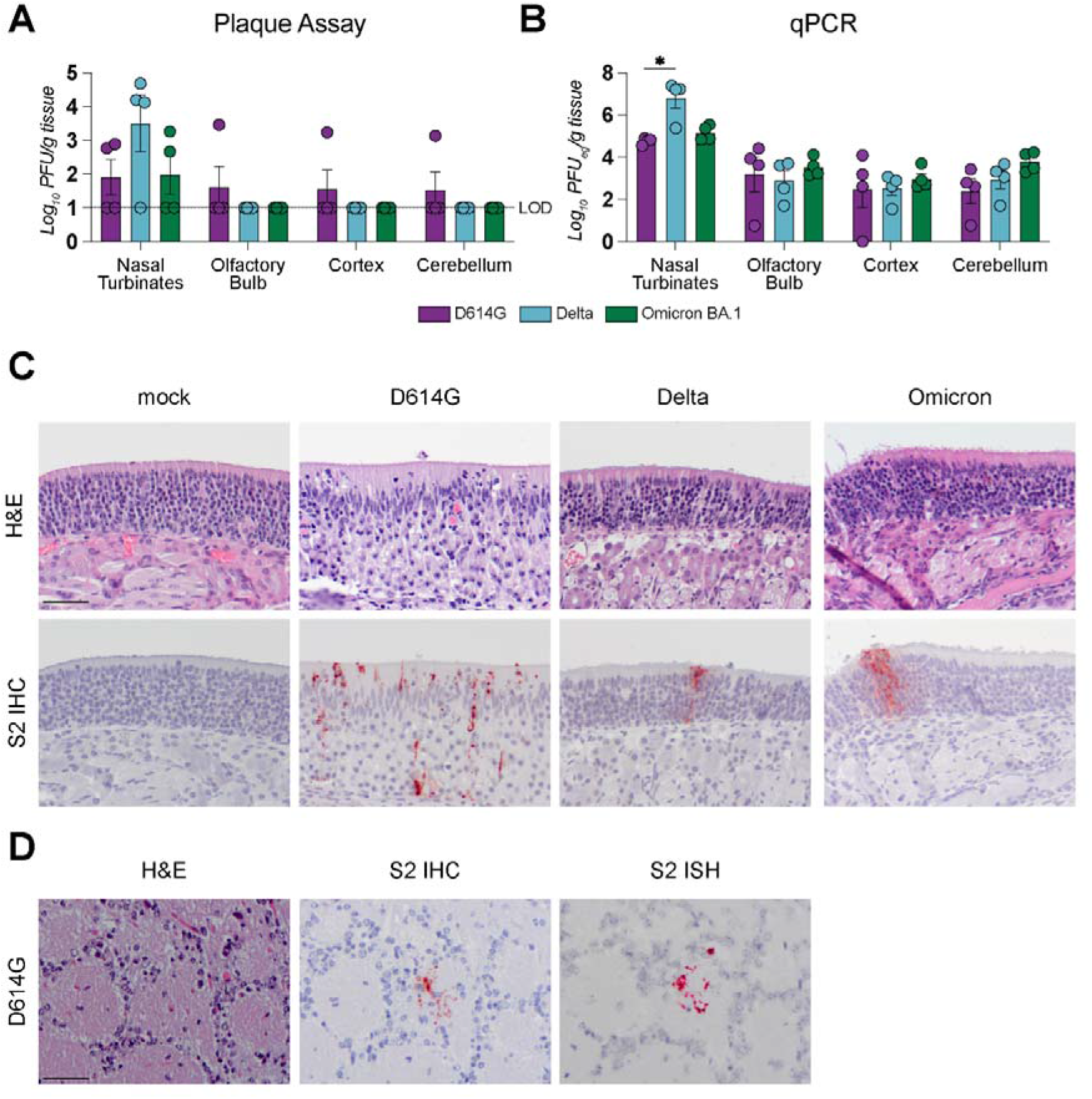
SARS-CoV-2 in nasal turbinates, olfactory bulb, cerebral cortex and cerebellum. Syrian golden hamsters were intranasally inoculated with 10^5^TCID_50_ D614G, 5.0×10^4^PFU Delta and Omicron BA.1 variant. Five days after inoculation, hamsters were sacrificed. Infectious virus titers (A) and viral copy RNA (B) in homogenates of nasal turbinates, olfactory bulb, cerebral cortex and cerebellum were quantified using plaque assay or reverse transcriptase quantitate PCR (RT-qPCR) respectively. Statistical significance was calculated with a Two-Way analysis of variance (ANOVA) with a Dunnett’s *posthoc* test. Averaged values of four individual animals per infection group were compared with all every other averaged value of four infected animals. Asterisks indicate statistical significance*, P<0.05. LOD, limit of detection (C) Hematoxylin and eosin (H&E) staining and SARS-CoV-2 nucleoprotein detected using immunohistochemistry (S2 IHC) in the olfactory mucosa of hamsters five days post exposure. (D) H&E staining SARS-CoV-2 nucleoprotein (S2 IHC) and SARS-CoV-2 viral RNA (S2 in-situ hybridization ISH) in the glomerular layer of the olfactory bulb in D614G inoculated hamsters.

Next, we determined the presence of virus, viral RNA and virus antigens in the olfactory bulb, cerebral cortex and cerebellum. In the olfactory bulb, infectious virus could be detected in one out of four D614G inoculated animals and in no animals being infected with the Delta or Omicron BA.1 variant (Figure 1A). However, similar levels of viral RNA were detected in the olfactory bulb in all groups (Figure 1B). Virus antigen, detected by IHC, was detected in the olfactory bulb of three out of four D614G infected animals (Table S1). Based on tissue architecture, location and cellular characteristics, the viral antigen was present in periglomerular cells of the glomerular layer. Varying between hamsters, one individual cell to several small clusters of these periglomerular cells had antigen. Viral RNA, detected by in-situ hybridization (ISH), was detected at the same sites in serial sections of the glomerular layers, as where SARS-CoV-2 antigen was detected (Figure 1D). No histological lesions were seen in any of the olfactory bulbs.

In the cerebral cortex, infectious virus was only isolated from one out of four D614G inoculated hamsters. Although not significant, higher viral titers were detected in D614G inoculated hamsters, compared to Delta and Omicron BA.1 inoculated hamsters. However, there was no evidence for the presence of virus antigen or histological lesions in the cerebral cortex of any of the hamsters.

In the cerebellum, infectious virus could be isolated in one out of four D614G inoculated hamsters and from none of the Delta or Omicron BA.1 infected hamsters, although there were no differences in viral RNA levels among the different groups. Similar to the cerebral cortex, there was no evidence for the presence of virus antigens or histological lesions in the cerebellum of any of the hamsters.

### Prominent antiviral and inflammatory response in the olfactory bulbs of hamsters infected with D614G, but not in hamsters inoculated with Delta or Omicron BA.1

To analyze the antiviral response, we first examined the expression of type-I and type-III-interferon (IFN) response in the olfactory bulb, cerebral cortex and cerebellum by reverse transcriptase quantitative PCR (RT-qPCR). Within the olfactory bulb, interferon-β (*Ifnb)* and interferon-λ (*Ifnl)* mRNA were upregulated in hamsters infected with D614G, but not in hamsters inoculated with the Delta or Omicron BA.1 variant (Figure 2A). Similarly, several canonical interferon-stimulated genes (ISGs) such as interferon-induced GTP-binding protein *Mx2,* interferon stimulated gene 15 (*Isg15)*, signal transducer and activator of transcription 1 (*Stat1)*, interferon regulatory protein 7 (*Irf)* were upregulated in the olfactory bulbs of the D614G hamsters but not in Delta or Omicron BA.1 inoculated hamsters (Figure 2A and Figure S2A). Induction of IFNs and ISGs were restricted to the olfactory bulb and could not be observed in the cerebral cortex or cerebellum of the hamsters (Figure 2B and 2C, Figure S2B and S2C).

**Figure 2.**
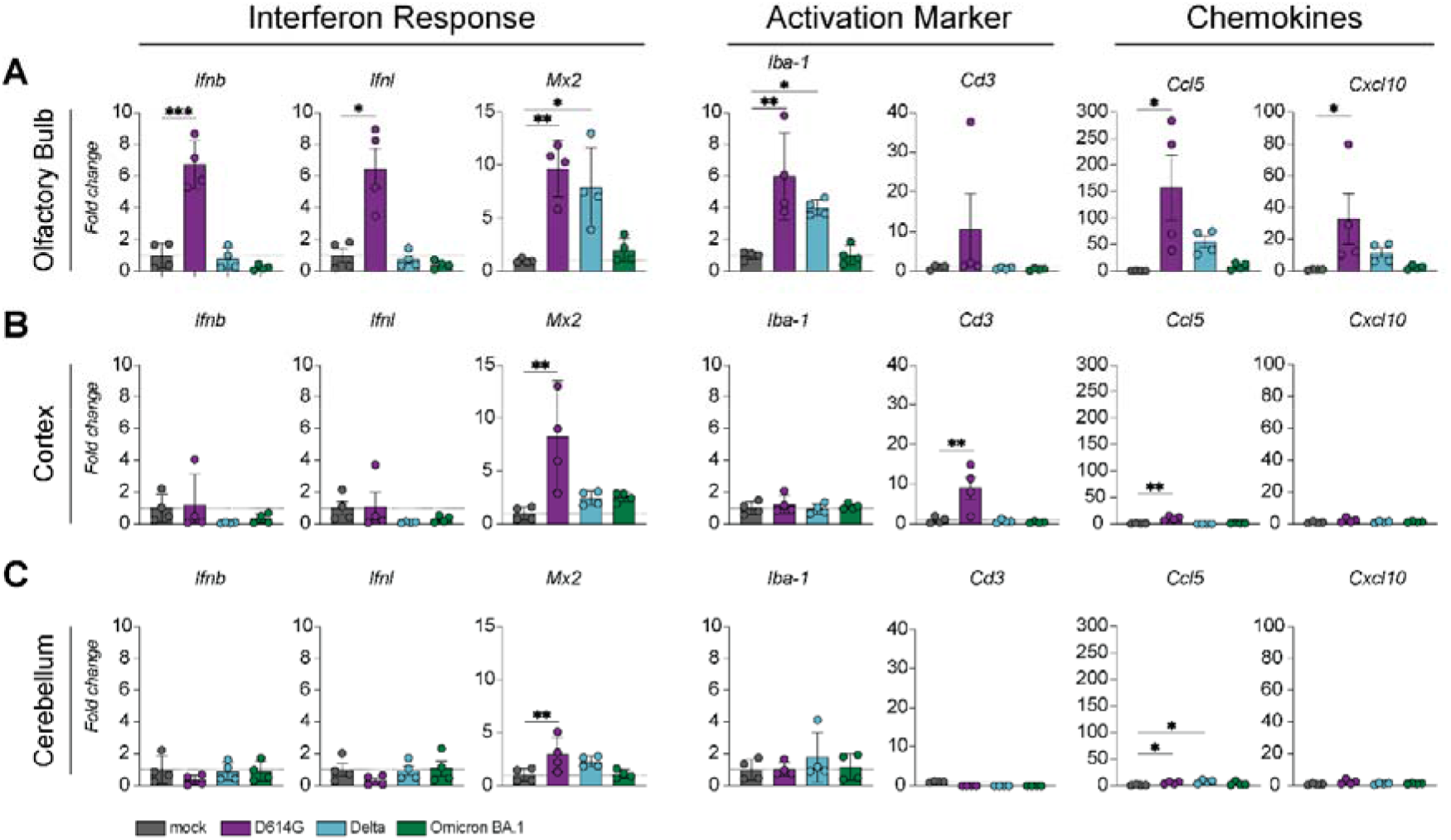
Antiviral and inflammatory responses in the olfactory bulb, cerebral cortex and cerebellum of SARS-CoV-2 infected hamsters. Expression of several genes important for interferon responses, for recruitment of microglia/macrophages and T-cells and chemokines were determined with reverse transcriptase quantitative PCR in the (A) olfactory bulb, (B) cerebral cortex and (C) cerebellum. The data displayed represent four animals per group. Statistical significance was calculated with a One-Way analysis of variance (ANOVA) with a Dunnett’s *posthoc* test. Averaged values of four individual animals per infection group were compared to values of four mock treated animals. Asterisks indicate statistical significance*, P<0.05, **, P<0.01, ***P<0.001, ****P>0.0001). Ifnb, interferon-β; Ifnl, interferon-λ; Mx2, interferon-induced GTP-binding protein; Aif1/Iba-1 allograft inflammatory factor 1 *Aif1* encoding the protein ionized calcium-binding adapter molecule 1 (IBA-1); Cd3, cluster of differentiation*; Cxcl10,* C-X-C motif chemokine 10; CCL5 C-C motif chemokine ligand 5 (*Ccl5*).

Next, we investigated the inflammatory response in the olfactory bulb, cerebral cortex and cerebellum. A significant increase of allograft inflammatory factor 1 *Aif1* mRNA, encoding the protein ionized calcium-binding adapter molecule 1 (IBA-1), was detected in the olfactory bulbs of hamsters inoculated with D614G or Delta, but not in the hamsters inoculated with the Omicron BA.1 variant (Figure 2B). The T cell associated gene cluster of differentiation 3 (*Cd3)* was not upregulated in the olfactory bulb of any of the groups. In the olfactory bulb of the D614G infected hamsters, but not in hamsters inoculated with Delta or Omicron BA.1, a significant increase of the chemokines C-X-C motif chemokine 10 (*Cxcl10)* and C-C motif chemokine ligand 5 (*Ccl5)* mRNA *was detected.* In the cerebral cortex, *Cd3* expression was only upregulated in D614G inoculated hamsters. In the cerebellum, there was no evidence for the induction of ISGs, proinflammatory cytokines and recruitment markers for macrophages/microglia (Figure 2B and Figure S2B)

Together, these data suggest that antiviral and inflammatory responses are predominantly located in the olfactory bulb five days post SARS-CoV-2 inoculation. This response was most prominent in hamsters inoculated with D614G. In the Delta infected hamsters we detected an upregulation of the ISG *Mx2* and the inflammatory marker *Iba-1* in the olfactory bulb. No antiviral or inflammatory response was detected in the olfactory bulb of hamsters infected with the Omicron BA.1 variant.

### Recruitment and activation of microglia/macrophages in the olfactory bulb of D614G- but not in Delta- or Omicron BA.1 infected hamsters

In order to confirm the increase of IBA-1 expression in the olfactory bulb of hamsters infected with D614G, we analyzed IBA-1 by IHC in the different layers of the olfactory bulbs of the D614G, Delta or Omicron BA.1 inoculated hamsters (Figure S2). In all hamster brains, including those of mock infected animals, we detected IBA-1 expression throughout the different layers of the olfactory bulb (Figure 3A and Figure S3). In all SARS-CoV-2 inoculated hamsters, the number as well as the size of IBA-1 expressing microglia increased. This was more apparent in hamsters inoculated with D614G, and less for Delta or Omicron BA.1 inoculated hamsters. To quantify this change, the number of IBA-1^+^ cells in the glomerular and the granule cell layer of the olfactory bulb were counted. An increase of IBA-1^+^ cells was observed in the glomerular layer of D614G infected hamsters, but not in Delta or Omicron BA.1 inoculated hamsters (Figure 3B). In the granule cell layer, IBA-1^+^ cells increased in the D614G hamsters, but not in the Delta and Omicron BA.1 hamsters. IBA-1^+^ cells clustered more frequently surrounding small blood vessels throughout all layers in D614G inoculated hamsters without any evidence for histological changes. (Figure S4). In accordance with *Cd3* mRNA expression, we did not observe and increase of CD3^+^ cells in any of the hamsters (Figure 3C).

**Figure 3.**
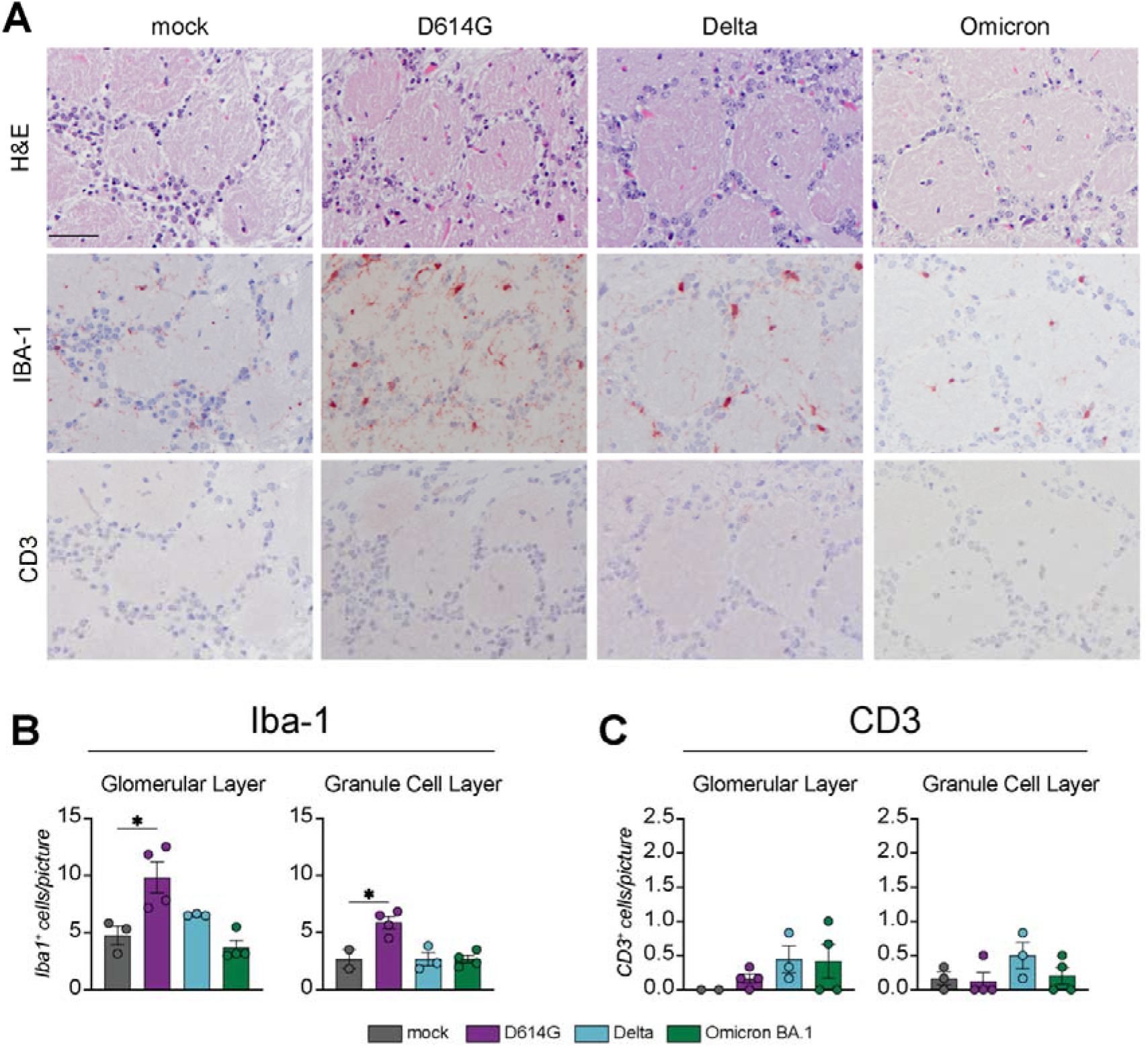
D614G infection but not Delta or Omicron BA.1 infection increases the number of Iba-1^+^ microglia/macrophages in the olfactory bulb. (A) Hematoxylin and eosin (H&E) staining of the glomerular layer in the olfactory bulb. Detection of Iba1^+^ cells and CD3^+^ cells in the glomerular layer of the olfactory bulb. Counting of (B) Iba-1^+^ cells and (C) CD3^+^ cells in the glomerular layer and the granular cell layer of the olfactory bulb. Statistical significance was calculated with a One-Way analysis of variance (ANOVA) with a Dunnett’s *posthoc* test. Averaged values of four individual animals per infection group were compared to values of four mock treated animals. Asterisks indicate statistical significance*, P<0.05)

## Discussion

Our study reveals that there are differences in the neuroinvasiveness and neurovirulence among the ancestral D614G and the Delta and Omicron BA.1 SARS-CoV-2 variants in the hamster model in the acute phase of the infection. D614G appears to be neuroinvasive, entering the CNS via the olfactory nerve, as previously observed in other studies (Frere et al., 2022; Imai et al., 2020; de Melo et al., 2021; Zazhytska et al., 2022). In contrast, although viral RNA was detected in the olfactory bulbs by RT-qPCR in Delta and Omicron BA.1 inoculated hamsters, this could not be confirmed by the detection of viral proteins or RNA by IHC and ISH respectively. This suggests a reduced neuroinvasive potential of these variants compared to the ancestral D614G variant, although we cannot exclude the possibility of neuroinvasion by the Delta and Omicron BA.1 variant at earlier timepoints after inoculation. As virus antigen and virus induced lesions were more abundant in the olfactory mucosa of D614G inoculated hamsters compared to the Delta or Omicron BA.1 inoculated hamsters, this might indicate that there is an association between the replication efficiency in the olfactory mucosa with the detection of virus antigen and viral RNA in the olfactory bulb. This pattern is also observed for influenza A viruses, where highly pathogenic H5N1 virus replicates efficiently in the olfactory mucosa and spreads to the CNS via the olfactory nerve (van den Brand et al., 2012; Schrauwen et al., 2012; Siegers et al., 2016), whereas, seasonal and pandemic influenza viruses replicate less efficient in the olfactory mucosa, and do not spread— at least not efficiently—to the CNS via the olfactory nerve (van den Brand et al., 2012; De Wit et al., 2018).

D614G appeared to be more neurovirulent than the Delta and Omicron BA.1 variant in the hamster models. The observed neurovirulence associated with D614G infection included antiviral and inflammatory responses in the olfactory bulb and to a lesser extent in the cerebral cortex, which fits with previous findings (Frere et al., 2022). On the contrary, infection with Delta or Omicron BA.1 variant hardly induced neuroinflammation in the olfactory bulb, cerebral cortex or cerebellum in this study. These observations fit with the fact that anosmia was observed more frequently in humans after infection with viruses containing the D614G mutation early in the pandemic (Von Bartheld et al., 2021). In addition, recent findings suggest that anosmia occurs less frequently after infection with Omicron BA.1, although these findings need to be validated in other studies (Maisa et al., 2022; Mutiawati et al., 2021). Whether microglial activation in the D614G inoculated hamsters contributes to anosmia or other neurological manifestations (Frere et al., 2022) or whether this is a protective response preventing subsequent virus spread throughout the CNS requires additional investigations.

The hamster model has been proven useful to study the pathogenesis of respiratory disease caused by SARS-CoV-2 infection for different variants (Bryche et al., 2020; Frere et al., 2022; de Melo et al., 2021; Rissmann et al., 2022; Sia et al., 2020; Zazhytska et al., 2022). This study suggests that the neuroinvasiveness and neurovirulence of different SARS-CoV-2 variants can also be studied in the hamster model. Here we show that infection with different SARS-CoV-2 variants reveals differences in neuroinvasive and neurovirulent potential in hamsters. As we only looked at one time point in the acute phase of the infection, future studies should reveal whether these findings can be extrapolated to the post-acute stage, and to the human situation.

## Material and Methods

### Cells

VeroE6 (ATCC CRL 1586) cells were maintained in Dulbecco’s modified Eagle’s medium (DMEM; Lonza,) supplemented with 10% fetal calf serum (FCS; Sigma-Aldrich,), 10◻mM HEPES, 1.5◻mg/ml sodium bicarbonate, 100◻IU/ml penicillin (Lonza,), and 100◻μg/ml streptomycin (Lonza). Calu-3 cells were cultured in Opti-MEM I (1) + GlutaMAX (Gibco) supplemented with 10% FBS, penicillin (100 IU/mL), and streptomycin (100 IU/mL). Both cells were grown at 37°C in a humidified CO2◻incubator and routinely tested for mycoplasma.

### Viruses

The SARS-CoV-2 D614G isolate (isolate BetaCoV/Munich/BavPat1/2020; European Virus Archive Global no. 026V-03883; kindly provided by C. Drosten) was propagated to passage three on Vero E6 cells in Opti-MEM I (1X) + GlutaMAX (Gibco), supplemented with penicillin (10,000 IU/mL) and streptomycin (10,000 IU/mL). Sequencing of the virus stock did not reveal major substitution as previously described in *Kutter et. al.* (Kutter et al., 2021). The SARS-CoV-2 variants Delta (B.1.617.2) and Omicron BA.1 (B.1.1.529) were propagated to passage three on Calu-3 cells in Advanced DMEM/F12 (Gibco), supplemented with HEPES, Glutamax, penicillin (100 IU/mL) and streptomycin (100 IU/mL) (AdDF+++). The Delta and Omicron BA.1 sequences are available on GenBank under accession numbers OM287123 and OM287553, respectively. All three viruses were grown◻at 37°C in a humidified CO2◻incubator. Infections were performed at a multiplicity of infection (MOI) of 0.01 and virus was harvested after 72 hours or at the peak of replication. The culture supernatant stored at −80°C. Virus titers were determined by plaque assay as described below.

All work was performed in a Class II Biosafety Cabinet under BSL-3 conditions at Erasmus Medical Center.

### Virus Titrations

Ten-fold serial diluted samples were added to monolayers of Calu-3 cells. Cells were incubated with inoculums at 37°C for 4 hours, washed once with PBS and then overlayed with 1.2% Avicel (FMC biopolymers) in Opti-MEM I (1X) + GlutaMAX for two days. Cells were fixed in 4% formalin for 20 minutes, permeabilized in 70% ice-cold ethanol and washed in PBS. Cells were incubated in 3% BSA (bovine serum albumin; Sigma) in PBS and stained with a rabbit anti-nucleocapsid antibody (Sino biological; 1:2000) in PBS containing 0.1% BSA, washed thrice in PBS, and stained with donkey anti-rabbit Alexa Fluor 488 (Invitrogen; 1:4000) in PBS containing 0.1% BSA. Cells were washed thrice in PBS and plates were scanned on the Amersham Typhoon Biomolecular Imager (channel Cy2; resolution 25 μm; GE Healthcare). All staining steps were performed at room temperature for one hour. Plaque assay analysis was performed using ImageQuant TL 8.2 software (GE Healthcare).

### Animals

#### Ethical Statement

Research involving animals was conducted in compliance with the Dutch legislation for the protection of animals used for scientific purposes (2014, implementing EU Directive 2010/63) and other relevant regulations. The licensed establishment where this research was conducted (Erasmus MC) has an approved OLAW Assurance #A5051-01. Research was conducted under a project license from the Dutch competent authority and the study protocol (#17-4312) was approved by the institutional Animal Welfare Body. Animals were housed in groups of 2 animals in filter top cages (T3, Techniplast), in Class III isolators allowing social interactions, under controlled conditions of humidity, temperature and light (12-hour light/12-hour dark cycles). Food and water were available ad libitum. Animals were cared for and monitored (pre- and post-infection) daily by qualified personnel. All animals were allowed to acclimatize to husbandry for at least 7 days. For unbiased experiments, all animals were randomly assigned to experimental groups. The animals were anesthetized (3-5% isoflurane) for all invasive procedures. Hamsters were euthanized by cardiac puncture under isoflurane anesthesia and cervical dislocation.

#### Animals and experimental setup

Female Syrian golden hamsters (*Mesocricetus auratus*; 6 weeks old; Janvier, France) were handled in an ABSL-3 biocontainment laboratory. Groups of animals (n = 4) were inoculated intranasally with 1 × 10^5^ TCID_50_ D614G, 5 × 10^4^ PFU of Omicron BA.1 or Delta variants of SARS-CoV-2 or PBS (mock; n = 4) in a total volume of 100μl per animal. On 5 dpi, infected animals were euthanized and the respiratory tract (nasal turbinate), as well as the CNS (olfactory bulb, cerebral cortex, cerebellum) was sampled for quantification of viral and genomic load, as well as for histopathology. Mock infected animals were euthanized 14 days post mock infection. Samples for histopathological analyses were fixed in 10% formalin for 2 weeks, after which tissues were paraffin-embedded

### Immunohistochemistry

For detection of SARS-CoV-2, Iba-1 and CD3 antigen in tissues of olfactory bulb and nose of hamsters, 3 μm formalin-fixed, paraffin-embedded sections were deparaffinized, rehydrated and pre-treated by boiling for 15 min in citric acid buffer pH 6.0 (SARS-CoV-2) or TRIS-EDTA pH 9.0 (Iba-1 and CD3). Endogenous peroxidase was blocked with 3% hydrogen peroxide for 10min at RT, after which slides were briefly washed with phosphate-buffered saline/0.05% Tween 20. Sections for SARS-CoV-2 IHC were blocked with 10% goat serum (X0907, DAKO, Agilent Technologies Netherlands B.V) for 30 min at RT. Slides were then incubated with a rabbit polyclonal antibody against SARS-CoV/SARS-CoV-2-nucleoprotein (40143-T62, Sino Biological, Pennsylvania, USA) (1:1000), mouse CD3 (ab16669, Abcam, Cambridge, UK) (1:10, 20μg/ml), human Iba-1 (019-19741, Wako Pure Chemical Corporation, Osaka, Japan) (1:200; 2.5μg/ml)) or Rabbit IgG isotypecontrol (AB-105-C, R&D, UK) (1:200; 5μg/ml) in PBS/0,1% BSA for 1h at room temperature (RT) (Table 1). After washing, sections were incubated with peroxidase labeled goat-anti-Rabbit IgG (1:100) (P0448, DAKO, Agilent Technologies Netherlands B.V) in PBS/0,1% BSA for 1h at RT. Peroxidase activity was revealed by incubating slides in 3-amino-9-ethylcarbazole (AEC) (Sigma) for 10 minutes, resulting in bright red precipitate, followed by counterstaining with hematoxylin. A lung section from an experimentally SARS-CoV-2 inoculated hamster and a brain and spleen section from a mock inoculated hamster were used as positive control for SARS-CoV-2, Iba-1 and CD3 staining respectively.

**Table 1:**
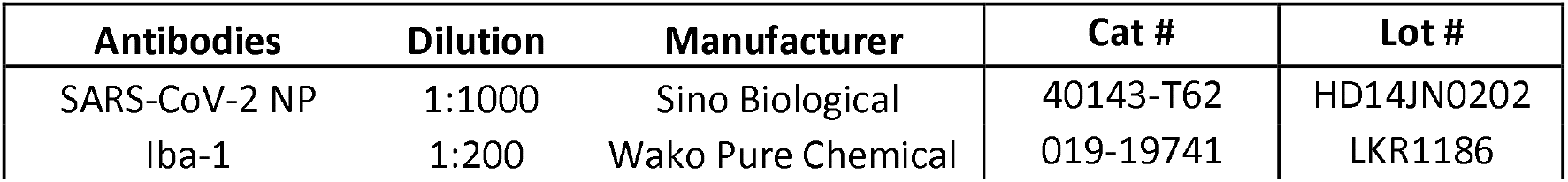
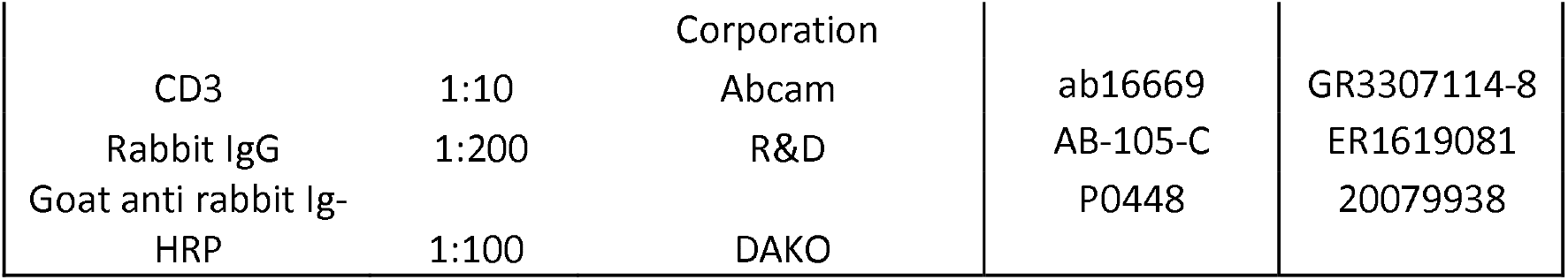
Antibodies used in this study

### SARS-CoV-2 In Situ Hybridization

BaseScope™ RNA probes were designed by Bio-Techne Ltd (Abingdon, UK) for BA-V-CoV-Wuhan-Nucleocapsid-3zz-st (846661). In situ hybridization was performed on formalin-fixed, paraffin-embedded consecutive sections from hamster olfactory bulb using BaseScope™ Reagent Kit v2–RED (323900) as described by the manufacturer.

### Counting of Microglia

From every animal three pictures of the granule cellular layer and the glomerular layer were taken. Counting of IBA-1^+^ cells was determined by manual counting by two independent blinded observes. Averages and Standard Deviation of the counting was plotted.

### Microscopy

A 10x air objective (Olympus) was used to select a region of interest with an Olympus BX51 microscope. Images were taken with a 200x magnification (20x air objective; Olympus) with CellSens software.

### Reverse transcriptase quantitative PCR

For the measurement of gene expression by RT-qPCR, total RNA was isolated as described previously in Lamers et al (Lamers et al., 2020). Briefly, 60 μL of sample was lysed in 90 μL of MagnaPure LC Lysis buffer (Roche) followed by a 30-minute incubation with 50 μL Agencourt AMPure XP beads (Beckman Coulter). Beads were washed thrice with 70% ethanol on a DynaMag-96 magnet (Invitrogen) and eluted in

50 μL diethylpyrocarbonate treated water. In total 500 ng of RNA were reverse transcribed with SuperScript™ IV Reverse Transcriptase using Random Hexamer Primers according to the manufacturers protocol (Promega). Subsequently, gene expression was determined with SYBR GREEN PCR Mastermix (Applied Biosystems) according to the manufacturers protocol on a 7500 Real Time PCR Cycler (Applied Biosystems) with gene specific primers listed in Table S2. Relative expression values were calculated with the 2^−ΔΔCT^ method and normalized to the average CT values of the housekeeping genes *Gapdh* and *Rpl18*. For quantification of SARS-Cov-2 specific RNA, a RT-qPCR targeting the E gene of SARS-CoV-2 was used as previously reported in Corman et al (Corman et al., 2020) and Ct values were compared to a standard curve derived from a titrated D614G virus stock.

### Statistical analysis

Statistical differences between experimental groups were determined as described in the figure legends. P values of ≤0.05 were considered significant. Graphs and statistical tests were made with GraphPad Prism version 9. Figures were prepared with Adobe Illustrator CC2019, Adobe Photoshop CC2019 and Biorender.

## Acknowledgments

We thank I. Visser, E. Marshall, D. Noack, D. van Eck-Schipper, J.M. Fentener van Vlissingen, Ingeborg van Middelkoop, Rianne Stam, Vincent Duiverman, Lars Vermaat and Vincent Vaes for assistance with the animal studies. This work was funded in part by a fellowship from the Netherlands Organization for Scientific Research (VIDI contract 91718308) and a EUR fellowship to D.V.R. and NIH contract SAVE and to BR and MPGK.

## Conflict of Interest

The authors declare no conflict of interest

## Notes

### Competing Interest Statement

The authors have declared no competing interest.

