## Supplement Material for "Differences in neuroinflammation in the olfactory bulb between D614G, Delta and Omicron BA.1 SARS-CoV-2 variants in the hamster model"

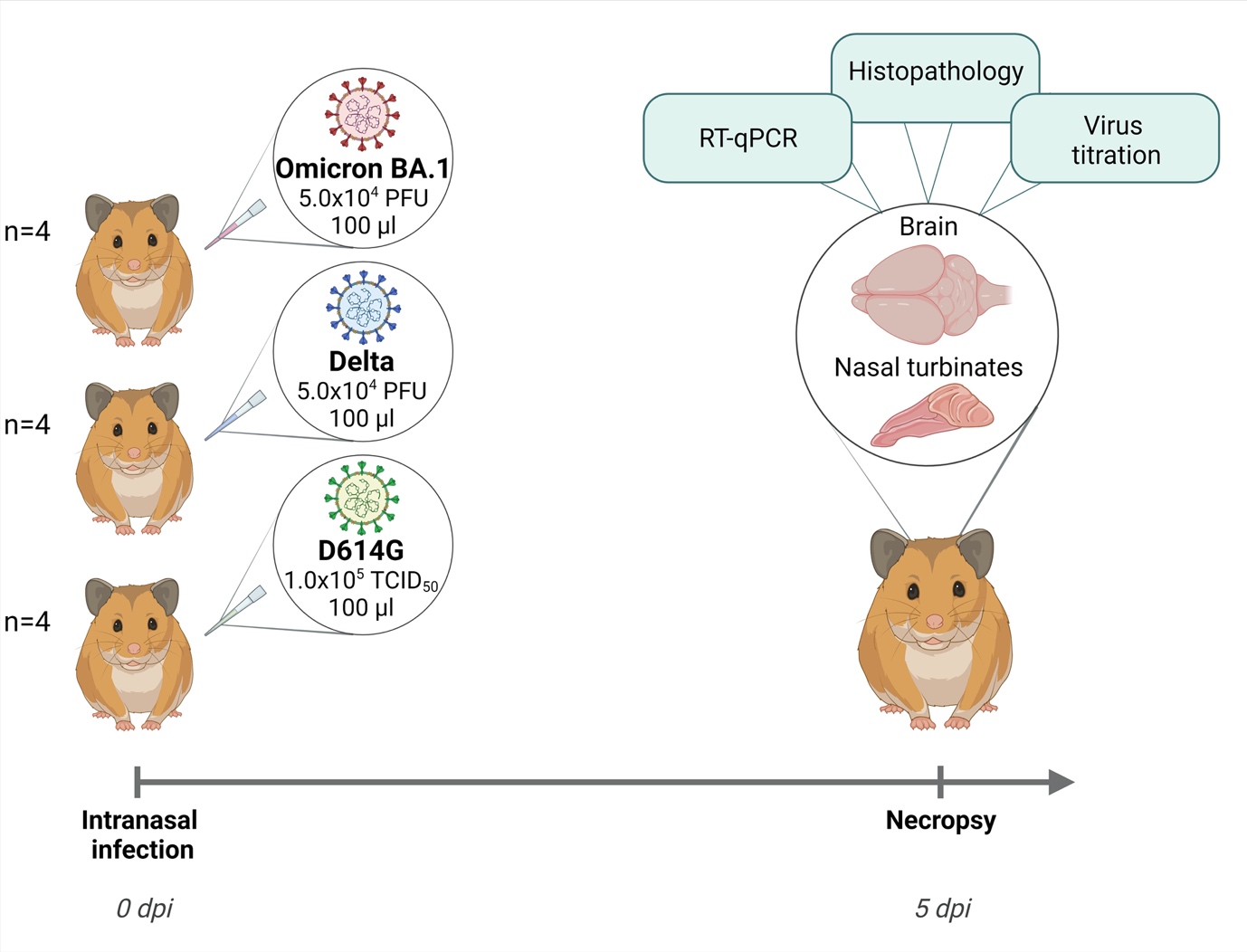

**Figure S1. Experimental set-up of the hamster experiment.** Syrian hamsters were intranasally inoculated with 10^5^TCID_50_ D614G, 5.0x10^4^PFU Delta and Omicron BA.1 variant. Five days after inoculation hamsters were sacrificed. Nasal turbinates, olfactory bulb, cortex and cerebellum were collected and subjected to virus titration, real-time quantitative PCR and histopathology. Figure created with Biorender.com.

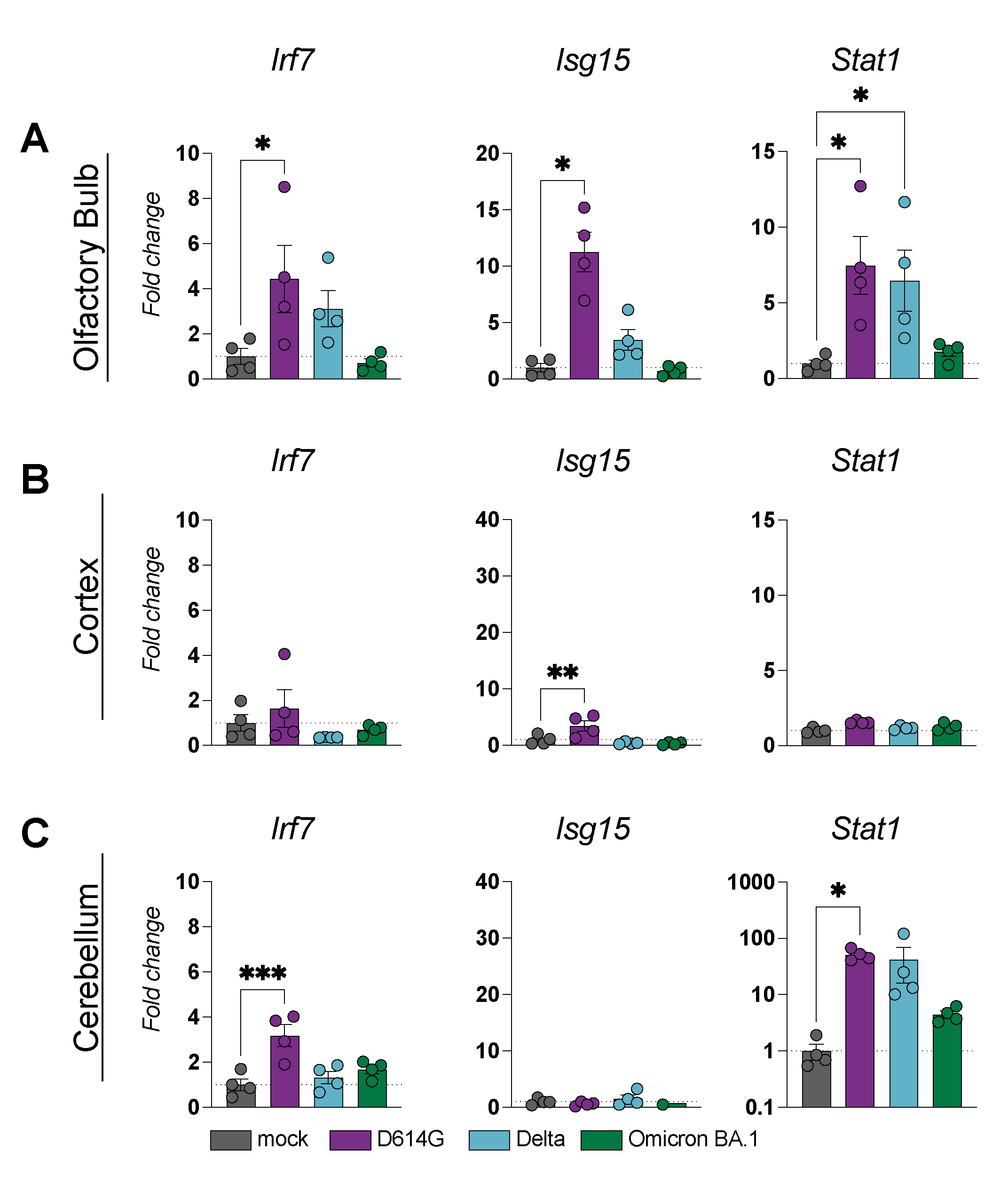

**Figure S2. Expression of interferon stimulated genes in the olfactory bulb, cortex and cerebellum of D614G, Delta and Omicron BA.1 infected hamster.** Expression level of interferon stimulated genes interferon regulatory protein 7 (*Irf7*), interferon stimulated gene 15 (*Isg15*) and signal transducer and activator of transcription 1 (*Stat1*) were determined with quantitative real-time PCR in the (A) olfactory bulb (B) cortex and (C) cerebellum. The data displayed represent four animals per group. Statistical significance was calculated with a One-Way analysis of variance (ANOVA) with a Dunnett’s *posthoc* test. Averaged values of four individual animals per infection group were compared to values of four mock treated animals. Asterisks indicate statistical significance*, P<0.05, **, P<0.01, ***P<0.001, ****P>0.0001). Irf7, interferon regulatory protein 7; Isg15, interferon stimulated gene 15; Stat1, signal transducer and activator of transcription 1

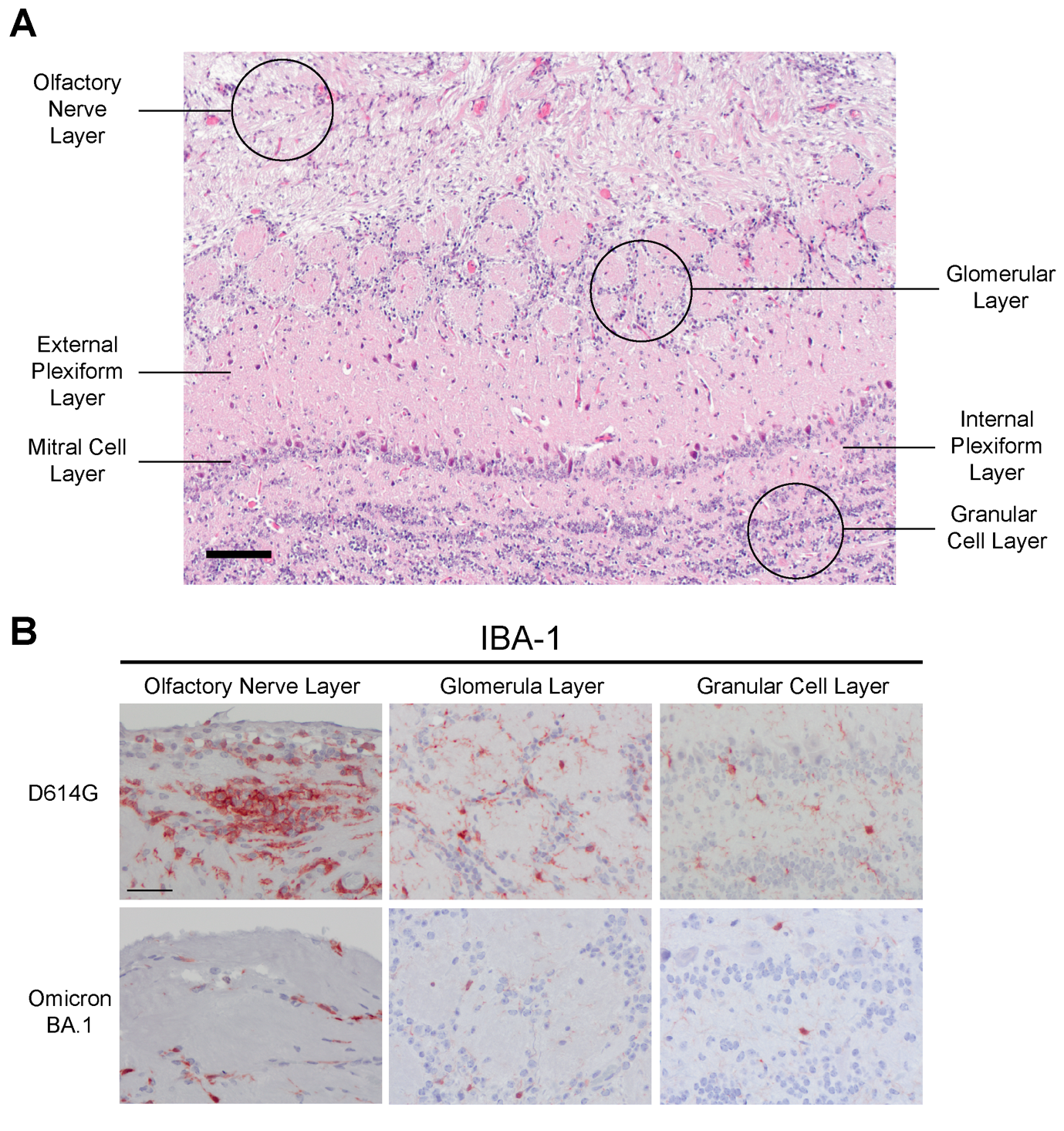

**Figure S2. Histological Overview of the olfactory bulb and the sampled regions.** (A) Overview of the olfactory bulb and its different layers. In the black circles, anatomical locations were highlighted which were used for analysis. (B) Comparison of ionized calcium-binding adapter molecule 1 (IBA-1) staining between D614G and Omicron BA.1 infected hamster in the olfactory nerve layer, glomerular layer and granular cell layer. Scale bars = 50µm.

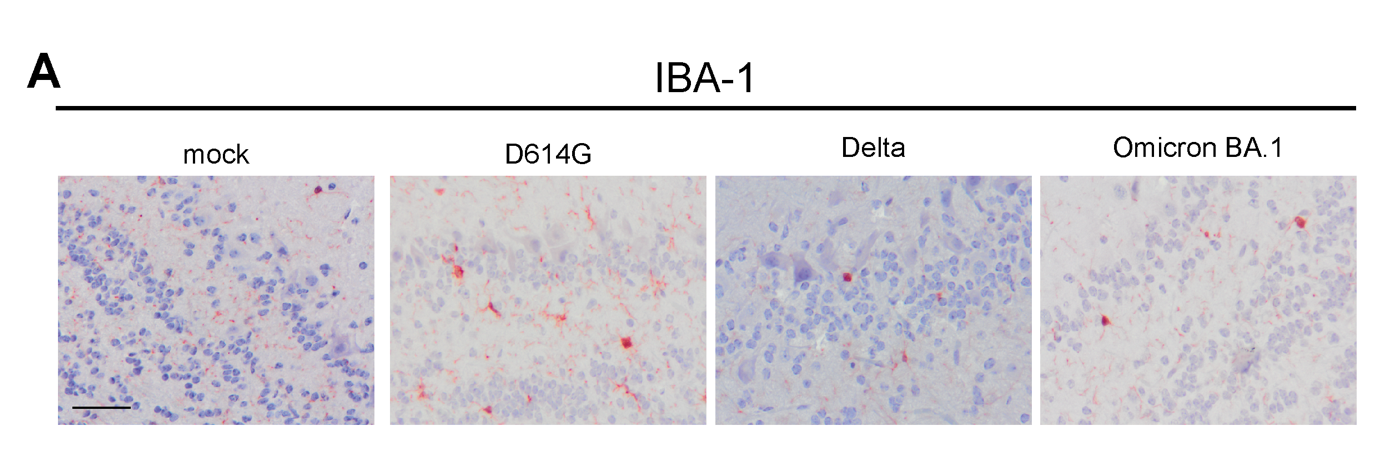

**Figure S3. Activation of microglia in the granular cell layer in the olfactory bulb.** (A) Detection of ionized calcium-binding adapter molecule 1 (IBA-1)^+^ cells in the granular cell layer of the olfactory bulb with immunohistochemistry. Scale bar = 50µM.

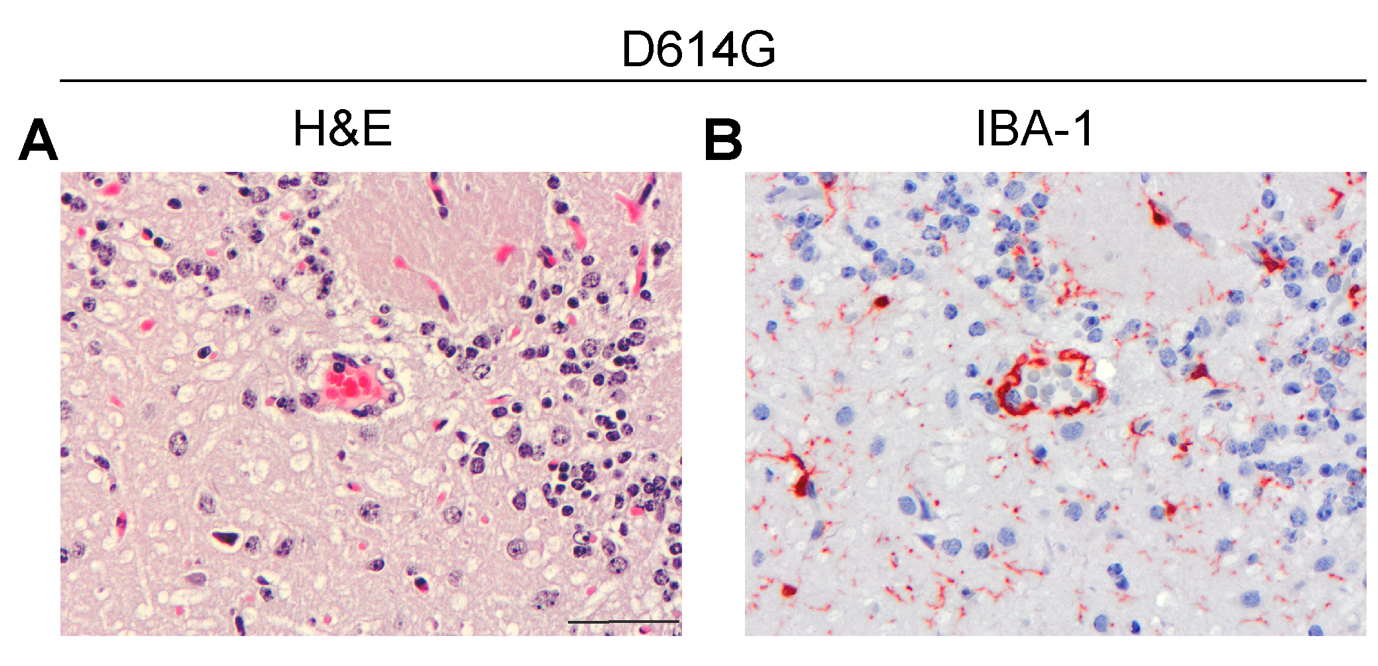

**Figure S4. Evidence for IBA-1^+^ cells at the level of the blood vessels**. (A) Hematoxylin and eosin (H&E) staining. (B) ionized calcium-binding adapter molecule 1 (IBA-1) immunohistochemistry indicates clustering of IBA-1^+^ cells around blood vessels in the granule cell layer of the olfactory bulb in D614G inoculated hamster. Scale bar = 50µM.

**Table S1. Pathological Scoring of olfactory mucosa and olfactory bulbs of infected hamsters.**

|  | | **Olfactory Mucosa** | | **Olfactory Bulb** | | |
| --- | --- | --- | --- | --- | --- | --- |
| **Group** | **Hamster** | **Inflammation** | **SARS-CoV-2 IHC** | **Inflammation** | **SARS-CoV-2 IHC** | **SARS-CoV-2 ISH** |
| **Mock** | 1 | None | - | None | - | - |
|  | 2 | None | - | None | - | - |
|  | 3 | None | - | None | - | - |
|  | 4 | None | - | None | - | - |
| **D614G** | 1 | N.A. | N.A. | None | + | + |
|  | 2 | Mild-moderate | + | None | - | - |
|  | 3 | N.A. | N.A. | None | + | + |
|  | 4 | Mild-moderate | + | None | + | + |
| **Delta** | 1 | Mild | + | None | - | - |
|  | 2 | Mild | + | N.A. | N.A. | N.A. |
|  | 3 | Mild-moderate | + | None | - | - |
|  | 4 | Mild | - | None | - | - |
| **Omicron** | 1 | Slight | + | None | - | - |
|  | 2 | Slight | + | None | - | - |
|  | 3 | None | + | None | - | - |
|  | 4 | Slight | + | None | - | - |

Sliding scale: none, slight, mild, moderate, marked, severe for inflammation
IHC – indicates no IHC positive cells, + indicated few-moderate number of IHC positive cells or ++ indicated many IHC positive cells
N.A. = not available

IHC, immunohistochemistry, ISH, in situ hybridization

**Table S2. Primers used for qPCR.**

| **Species** | **Genes** | **Sequence 5'-3'** | **Annealing °C** | **Amplicons** | **Ref** |
| --- | --- | --- | --- | --- | --- |
| *M.auratus* | ISG15_F | TCTATGAGGTCCGGCTGACA | 60 | 155 bp | [1] |
| *M.auratus* | ISG15_R | GCACTGGGGCTTTAGGTCAT |  |  |  |
| *M.auratus* | MX2_F | AGGCAGTGGTATTGTCACCAG | 60 | 187 bp | [1] |
| *M.auratus* | MX2_R | ATCACTGATCCCCAGCCCTAT |  |  |  |
| *M.auratus* | IRF7_F | ATTTCGGTCGCAGGGATCTG | 60 | 184 bp | [1] |
| *M.auratus* | IRF7_R | TGCAAGATAAAGCGTCCCGT |  |  |  |
| *M.auratus* | CXCL10_F | TCTGAGTGGGACTCAAGGAATC | 60 | 74 bp | [1] |
| *M.auratus* | CXCL10_R | CCGGTCGGTCATCGATTTTG |  |  |  |
| *M.auratus* | CCL5_F | TCTCCACAGCTGTCCTCACT | 58 | 198 bp | [1] |
| *M.auratus* | CCL5_R | TCCTTCGGGTGACAAAAACGA |  |  |  |
| *M.auratus* | IBA-1_F | GGGGAAAAGCCTTTGGACTG | 60 | 148 bp | [1] |
| *M.auratus* | IBA-1_R | TCAAACTCCATGTACTTCTTCTTGA |  |  |  |
| *M.auratus* | IFNb_F | CCATCATGACCAACAGGTGGA | 60 | 96 bp | [2] |
| *M.auratus* | IFNb_R | GTCTGGCCTCAAGTTCCTCG |  |  |  |
| *M.auratus* | CD3b_F | CTGGCTGCTTTTCTCCCTCG | 60 | 122 bp | [3] |
| *M.auratus* | CD3b_R | AACCATACTTTCGCCGTTCCC |  |  |  |
| *M.auratus* | IFNL_F | CCCACCAGATGCAAAGGATT | 60 | 109 bp | [4] |
| *M.auratus* | IFNL_R | CTTGAGCAGCCACTCTTCTATG |  |  |  |
| *M.auratus* | GAPDH_F | AACTTTGGCATTGTGGAAGG | 60 | 245 bp | [2] |
| *M.auratus* | GAPDH_R | CGACATGTGAGATCCACGA |  |  |  |
| *M.auratus* | RPL18_F | GTTTATGAGTCGCACTAACCG | 60 | 80 bp | [5] |
| *M.auratus* | RPL18_R | TGTTCTCTCGGCCAGGAA |  |  |  |
